# Comprehensive Analysis of CRISPR Array Repeat Mutations Reveals Subtype-Specific Patterns and Links to Spacer Dynamics

**DOI:** 10.1101/2025.04.02.646798

**Authors:** Alexander Mitrofanov, Chase L. Beisel, Franz Baumdicker, Omer S. Alkhnbashi, Rolf Backofen

**Author notes:** To whom correspondence should be addressed: Rolf Backofen, Omer S. Alkhnbashi.

## Abstract

CRISPR–Cas systems are adaptive immune mechanisms in bacteria and archaea that protect against invading genetic elements by integrating short fragments of foreign DNA into CRISPR arrays. These arrays consist of repetitive sequences interspersed with unique spacers, guiding Cas proteins to recognize and degrade matching nucleic acids. The integrity of these repeat sequences is crucial for the proper function of CRISPR–Cas systems, yet their mutational dynamics remain poorly understood.

In this study, we analyzed 56,343 CRISPR arrays across 25,628 diverse prokaryotic genomes to assess the mutation patterns in CRISPR array repeat sequences within and across different CRISPR subtypes. Our findings reveal, as expected to some extent, that mutation frequency is substantially higher in terminal repeat sequences compared to internal repeats consistently across system types. However, the mutation patterns exhibit an unexpected amount of variation among different CRISPR subtypes, suggesting that selective pressures and functional constraints shape repeat sequence evolution in distinct ways. Understanding these mutation dynamics provides insights into the stability and adaptability of CRISPR arrays across diverse bacterial and archaeal lineages. Additionally, we elucidate a novel relationship between repeat mutations and spacer dynamics, demonstrating that hotspots for terminal repeat mutations coincide with regions exhibiting spacer conservation. This observation corroborates recent findings by Fehrenbach et al. (2024) indicating that spacer deletions occur at a frequency 374 times greater than that of mutations and are significantly influenced by repeat misalignment. Our findings suggest that repeat mutations play a pivotal role in spacer retention or loss, or vice versa, thereby highlighting an evolutionary trade-off between the stability and adaptability of CRISPR arrays.

## Introduction

Clustered Regularly Interspaced Short Palindromic Repeats (CRISPR) and their associated CRISPR-associated (Cas) proteins constitute a widespread adaptive immune mechanism in archaea and bacteria, providing defense against invading genetic elements such as bacteriophages and conjugative plasmids. The CRISPR-Cas immune response is characterized by three critical stages: adaptation, processing, and interference. In the adaptation phase, short DNA segments derived from invaders are incorporated into the host genome as spacers, which are flanked by highly conserved repeat sequences, thereby forming a CRISPR array. Following this phase, the arrays are transcribed and processed into CRISPR RNAs (crRNAs), which subsequently guide Cas protein complexes to identify and, in many cases, cleave complementary nucleic acids during the interference phase (Lange et al., 2013; Alkhnbashi et al., 2014, 2016, 2021; Maier et al., 2017; Reimann et al., 2017; Stachler et al., 2020). CRISPR systems are currently classified into two major classes and seven types (Types I–VII) and over 30 subtypes and variants, each exhibiting unique effector protein compositions and functional mechanisms. Class 1 systems (e.g., Types I, III) employ multi-protein effector complexes, while Class 2 systems (e.g., Type II) use single, multidomain proteins like Cas9, enabling efficient genome editing applications. While proteins such as Cas1 and Cas2 (involved in spacer acquisition) are conserved across systems, others like Cas7 or Cas8 are more diverse, making their evolutionary analysis and identification challenging (Makarova et al., 2011, 2015, 2020; Vestergaard, Garrett and Shah, 2014; Koonin and Makarova, 2017; Makarova, Gao, et al., 2019; Makarova, Karamycheva, et al., 2019; Shah et al., 2019). The repeat sequences within CRISPR arrays are fundamental to all phases of the immune response. These direct repeats, which typically range from 24 to 47 base pairs in length, serve a critical role beyond mere structural components. They are essential for the recognition by Cas proteins, the processing of crRNA, and the acquisition of spacers. While repeat sequences are frequently characterized as conserved, recent research has revealed considerable sequence variability among them, both within individual CRISPR subtypes and across different types. This variability in repeat sequences may influence the efficacy of CRISPR functionality, particularly in relation to crRNA maturation and interference activities, where specific sequence motifs are vital for optimal immune defense (Kunin, Sorek and Hugenholtz, 2007; Lange et al., 2013; Charpentier et al., 2015). Importantly, mutations in repeat sequences reveal significant implications for both the evolution as well as the functionality of CRISPR systems. Certain mutations may emerge from errors during replication or the acquisition of spacers, while others may be influenced by positive or purifying selection, contingent upon their specific location and functional consequences. Recent findings indicate that repeat mutations are often non-randomly distributed, frequently accumulating at the terminal or leader-proximal regions of CRISPR arrays.

Repeat mutations are of particular interest due to their capability of indicating evolutionary signals. In more detail, differences in mutation patterns across CRISPR subtypes suggest that repeat sequences are subject to distinct selective pressures (Kunin, Sorek and Hugenholtz, 2007; Lange et al., 2013). Some subtypes may favor higher repeat stability to maintain the integrity of the CRISPR array, while others may tolerate or even benefit from mutations that enhance adaptability (van der Oost et al., 2014; Charpentier et al., 2015). These mutations may also contribute to the structural evolution of CRISPR loci, influencing how bacterial genomes respond to environmental pressures (Touchon and Rocha, 2010; Liao et al., 2022; Naeem and Alkhnbashi, 2023); (Touchon and Rocha, 2010; Westra *et al*., 2012; Staals *et al*., 2013)(Touchon and Rocha, 2010; Liao *et al*., 2022; Naeem and Alkhnbashi, 2023).

Beyond their evolutionary role, repeat mutations can have direct functional consequences on the efficiency and stability of CRISPR-Cas activity. Since repeat sequences serve as critical recognition elements for the processing and maturation of crRNAs, mutations within these sequences may impact spacer acquisition, crRNA formation, and interference with foreign DNA (Charpentier et al., 2015; Nickel et al., 2019; Stachler et al., 2020). In Type II CRISPR-Cas systems, for example, the Cas9 endonuclease relies on precise repeat sequences for proper crRNA processing (Deltcheva et al., 2011). Similarly, in Type I and III systems, repeat mutations may alter interactions with the Cascade complex or Cas10, respectively, affecting immune responses (Brouns et al., 2008; Hale et al., 2009, 2014; Makarova et al., 2011; Westra et al., 2012).

A specifically interesting aspect is that repeat mutation patterns vary across CRISPR subtypes, thus highlighting potential differences in their underlying mechanisms of spacer acquisition and interference (Makarova et al., 2020). Some subtypes may exhibit greater repeat sequence conservation to preserve CRISPR array stability, while others display higher mutation numbers that could provide flexibility in adapting to new threats (Staals et al., 2013, 2016). Understanding these subtype-specific variations could provide deeper insights into CRISPR-Cas evolution and functionality (Koonin and Makarova, 2019).

From an applied perspective, repeat mutations have significant implications for biotechnology and genome editing. The stability of CRISPR repeats is crucial in optimizing synthetic CRISPR systems for precise gene editing, ensuring robust guide RNA formation and minimizing off-target effects (Jinek et al., 2012). Additionally, engineered bacterial strains used in industrial and medical applications rely on stable CRISPR arrays for long-term phage resistance (Barrangou and Horvath, 2017). A deeper understanding of repeat mutation patterns and associated constraints can therefore enhance the design of more efficient CRISPR-based tools in synthetic biology and therapeutic applications (Hsu, Lander and Zhang, 2014; de la Fuente-Núñez and Lu, 2017; Abdelaal and Yazdani, 2020).

In this study, we conduct a comprehensive analysis of repeat mutations across diverse CRISPR-Cas subtypes, investigating their distribution, positional bias, and potential functional impact using a dataset across diverse prokaryotic lineages and leveraging robust bioinformatic tools for orientation and mutation analysis.

## Materials and methods

### Dataset acquisition

The dataset used in this study was constructed by integrating data from the CRISPRCasFinder (Couvin et al., 2018) database, a curated resource containing CRISPR arrays and associated Cas proteins identified across diverse prokaryotic genomes. To enhance the diversity and coverage of CRISPR-Cas system classifications, we further enriched our dataset with data from Makarova et al., which provides an updated evolutionary classification of CRISPR-Cas systems, including annotations of various types and subtypes (Makarova et al., 2011, 2015). The integration of these datasets resulted in a comprehensive and high-confidence collection of CRISPR-Cas system data, culminating in a total of 25628 unique genomes, ensuring a robust foundation for computational analysis and classification.

### Dataset processing

CRISPR array identification was performed using CRISPRCasFinder (Couvin et al., 2018) and CRISPRidentify (Mitrofanov *et al*., 2021), two established tools for detecting CRISPR arrays in genomic sequences. Both tools assign confidence levels to the detected arrays based on their likelihood of being true CRISPR structures. To ensure high reliability in our dataset, we applied stringent filtering criteria: only arrays with a confidence level of 4 were retained from the CRISPRCasFinder dataset. Similarly, from the CRISPRidentify dataset, only candidates classified as Bona-Fide and Possible were included. This approach minimized the inclusion of false positives while maintaining a broad and representative selection of CRISPR arrays for downstream analyses.

The identification and labeling of Cas genes were conducted using the Cas identification module of CRISPRCasFinder and CRISPRcasIdentifier (Padilha et al., 2020), both of which employ specialized algorithms to detect and classify Cas proteins within genomic sequences. Following the identification of Cas genes, each CRISPR array was assigned to its closest corresponding Cas locus. To ensure a biologically relevant association while minimizing erroneous links, a maximum distance threshold of 10,000 nucleotides was applied. Only arrays within this proximity to a detected Cas locus were considered associated, ensuring a robust and functionally meaningful mapping between CRISPR arrays and *cas* genes.

The orientation of CRISPR arrays was determined using three independent methods to enhance accuracy and reliability. CRISPRDetect (Naito et al., 2015), which is integrated within CRISPRCasFinder standalone version, was utilized as one approach for strand orientation prediction. Additionally, CRISPRStrand (Alkhnbashi et al., 2014), the internal orientation prediction module of CRISPRidentify (Mitrofanov et al., 2021), was employed to provide further validation. Lastly, CRISPRevoinator, an evolutionary-based tool built upon SpacerPlacer (Fehrenbach et al., 2024), was used to infer orientation based on evolutionary conservation and spacer sequence analysis. By leveraging these complementary methods, we ensured a robust and consistent prediction of CRISPR array orientation across the dataset.

## Results

### Potential Role of Repeat Mutations in Spacer Acquisition and Stability

The research conducted by Fehrenbach et al. (2024), which traces evolution of arrays purely based on their composition of evolutionarily conserved spacers, indicates that spacer deletions are more prevalent in the central regions of CRISPR arrays, while terminal spacers demonstrate a higher degree of conservation. Our examination of repeat mutation distributions of 56,343 CRISPR arrays supports this finding, revealing that terminal repeats are subject to a significantly greater frequency of mutations in comparison to internal repeats (See Table 1). This observation indicates that the stability of repeat sequences may be crucial in protecting adjacent spacers from deletion, thereby preserving the functionality of CRISPR arrays. It could also imply that a lower deletion rate of repeats at the array termini facilitates the accumulation of mutations at both ends. Both effects would show a similar result if they were independent events, however, they could even be mutually reinforcing. I.e., the requirement of a stable terminal repeat sequence might directly or indirectly lead to a lower deletion rate of terminal spacers. Moreover, our analysis identified that specific subtypes of CRISPR systems, notably II-C, display increased levels of repeat mutations at the leader-proximal end (See Table 1). This observation is consistent with previous research suggesting that the rates of spacer acquisition are influenced by variability in leader sequences. It is plausible that mutations occurring in the repeats located near the leader end may impact the recognition sites for the Cas1-Cas2 complex, thus affecting the efficiency of new spacer incorporation (Erdmann and Garrett, 2012; Yosef, Goren and Qimron, 2012; Díez-Villaseñor et al., 2013). Additionally, it has been proposed that repeat misalignment during replication may significantly contribute to spacer deletions (Garrett, 2021; Fehrenbach et al., 2024).

**Table 1.**
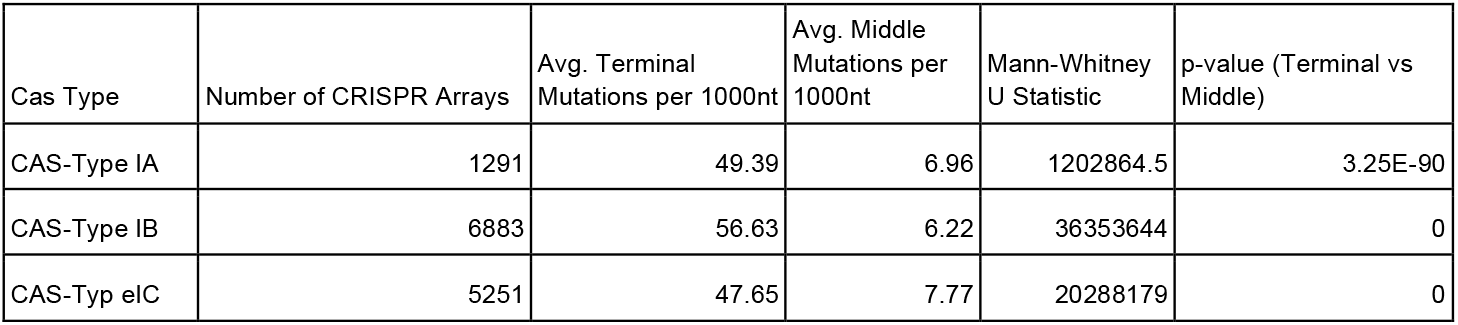
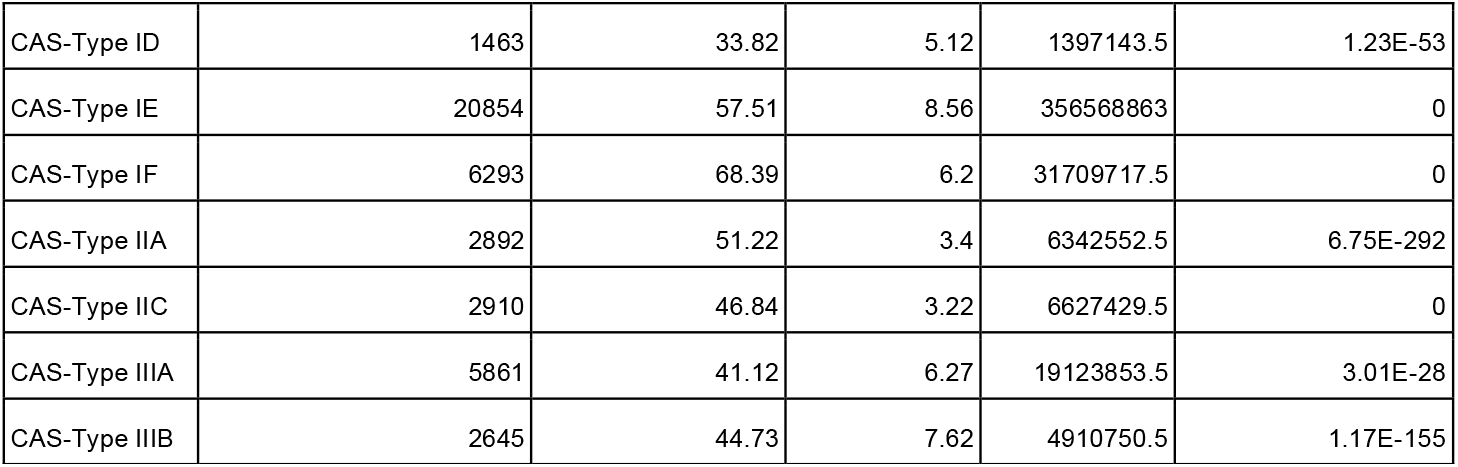
Overview of mutation frequencies (per 1000 nucleotides) across different CRISPR subtypes. The table highlights the distribution of mutations between terminal and middle repeat sequences. The data reveal substantial variation among CRISPR subtypes, with terminal repeats consistently showing significantly more mutations than middle repeats. Mann-Whitney tests conducted for each CRISPR subtype confirm that these differences are statistically significant.

| Cas Type | Number of CRISPR Arrays | Avg. Terminal Mutations per 1000nt | Avg. Middle Mutations per 1000nt | Mann-Whitney U Statistic | p-value (Terminal vs Middle) |
| --- | --- | --- | --- | --- | --- |
| CAS-Type IA | 1291 | 49.39 | 6.96 | 1202864.5 | 3.25E-90 |
| CAS-Type IB | 6883 | 56.63 | 6.22 | 36353644 | 0 |
| CAS-Typ eIC | 5251 | 47.65 | 7.77 | 20288179 | 0 |
| CAS-Type ID | 1463 | 33.82 | 5.12 | 1397143.5 | 1.23E-53 |
| CAS-Type IE | 20854 | 57.51 | 8.56 | 356568863 | 0 |
| CAS-Type IF | 6293 | 68.39 | 6.2 | 31709717.5 | 0 |
| CAS-Type IIA | 2892 | 51.22 | 3.4 | 6342552.5 | 6.75E-292 |
| CAS-Type IIC | 2910 | 46.84 | 3.22 | 6627429.5 | 0 |
| CAS-Type IIIA | 5861 | 41.12 | 6.27 | 19123853.5 | 3.01E-28 |
| CAS-Type IIIB | 2645 | 44.73 | 7.62 | 4910750.5 | 1.17E-155 |

Should repeat mutations compromise sequence homology to the other repeats in the array, they may modify the probability of spacer loss, most likely recombination-driven. Thus, it was shown that mutations in the last repeat could contribute to the lower spacer deletion rate towards the distal end of CRISPR arrays (Fehrenbach et al., 2024). This understanding may also clarify why certain CRISPR subtypes exhibit higher numbers of mutations at the array termini, potentially reflecting an evolutionary trade-off between adaptive flexibility and the maintenance of immune memory.

### Proximal/Terminal repeat mutations occur more often across all CRISPR subtypes

To investigate the distribution of mutations within CRISPR arrays, we analyzed the average number of mutations across the entire array. The objective was to determine whether mutations occur homogeneously throughout the array or if specific regions exhibit higher variability. Our results indicate that repeat mutations are not uniformly distributed but instead show a significantly higher frequency at both ends of the CRISPR arrays. This pattern was consistently observed across all CRISPR subtypes (see “Avg. Terminal Mutations per 1000 nt.” column in Table 1_)_, suggesting a potential evolutionary or mechanistic bias influencing the stability of repeats at the array termini.

The dataset provides a comprehensive analysis of mutation frequencies across different CRISPR subtypes, revealing distinct patterns in mutation distribution (see Table 1). The overall number of observed mutations per 1000 nucleotides varies significantly among CRISPR subtypes, ranging from the lowest in CAS-IIC (9.01 per 1000 nt) and CAS-ID (9.22 per 1000 nt) to the highest in CAS-IE (15.95 per 1000 nt) and CAS-IF (15.15 per 1000 nt). A notable trend across all types is the markedly higher frequency of terminal mutations compared to middle mutations (columns 4 and 5 in Table 1), suggesting a bias toward sequence alterations at the ends of the targeted regions. CAS-IF exhibits the highest number of terminal mutations (68.39 per 1000 nt), while middle mutations are relatively stable across CRISPR subtypes, generally ranging between 3 and 8 per 1000 nt. Statistical analysis through the Mann-Whitney U Statistic further confirms significant differences in mutation distributions, with p-values approaching zero in most cases, highlighting strong variation among CRISPR subtypes.

### The number of mutations in Proximal/Terminal repeats is different for different CRISPR subtypes

Following the observation of a predominant number of mutations in the terminal repeats, we further examined whether the occurrence of mutations varied among different CRISPR subtypes. To assess this, we analyzed the frequency of repeat mutations across distinct Cas system classifications. Our findings indicate that mutation patterns differ significantly among CRISPR subtypes (See Figure 1), suggesting that the evolutionary dynamics and stability of CRISPR arrays may be influenced by the specific Cas protein machinery associated with each system (See Figure 1).

**Figure 1.**
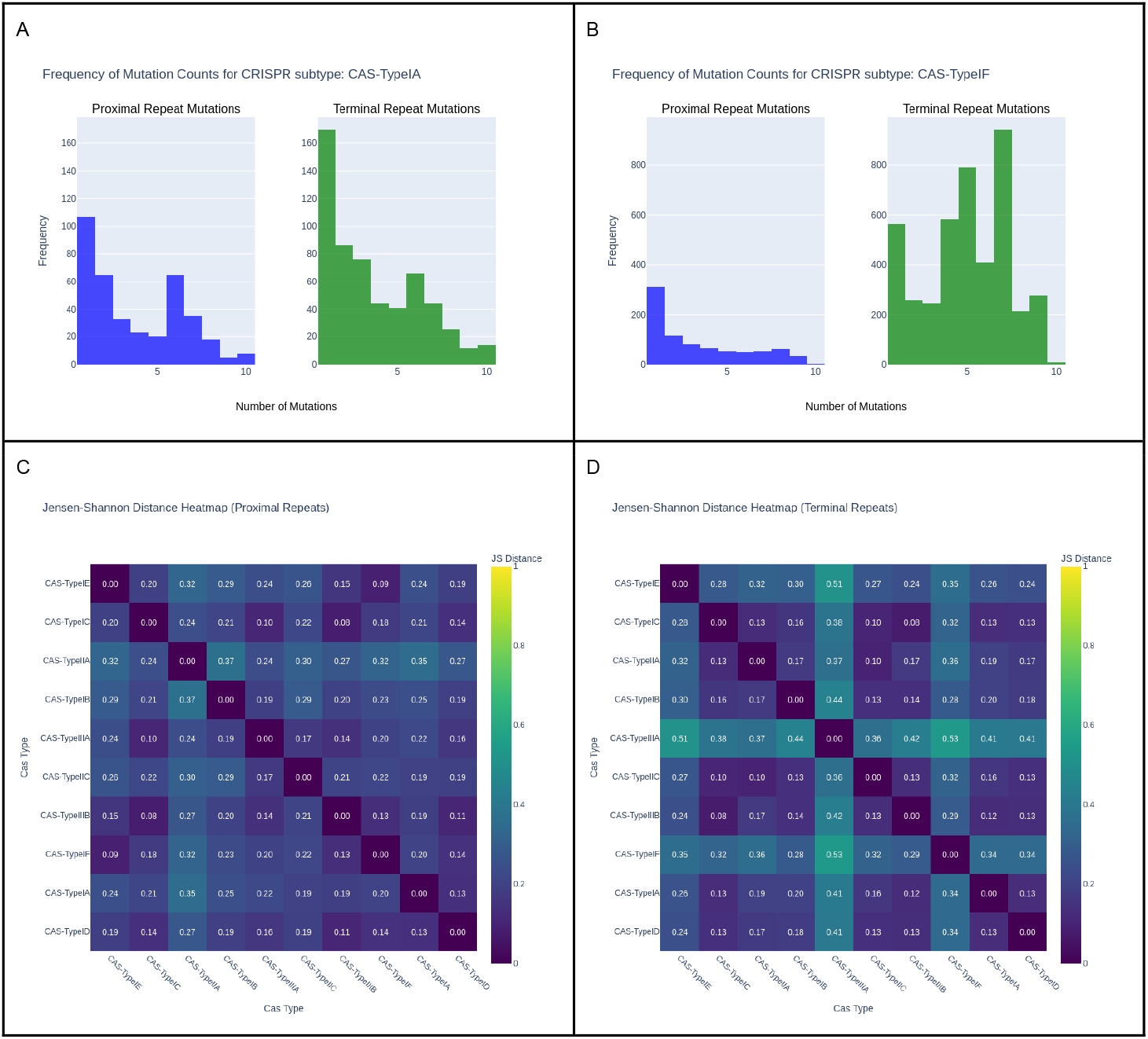
Mutation frequencies of proximal and terminal repeats across different CRISPR subtypes. The Figure shows histograms of mutation frequencies of the proximal and terminal repeats for CRISPR arrays of Type IA (panel A) and Type IF (Panel B). The histograms clearly show the difference in mutation frequency clearly indicating that type IF CRISPR arrays terminal repeats tend to have significantly higher mutated terminal repeat sequences.

### Mutation position distribution through the repeat sequence shows different patterns for different CRISPR subtypes

In the previous section, we investigated the number of mutations in repeats and their relation to the position of the repeat within the array (terminal or middle). In this section, we want to investigate in more detail where the repeat mutations occur. Thus, to examine the positional distribution of mutations within CRISPR repeats across different CRISPR subtypes, we analyzed how mutations were distributed along the repeat sequence. Given that repeat lengths vary across different CRISPR arrays, direct comparison was not feasible. To account for this variation, we applied length normalization, standardizing all repeats to a 100-bin interval. Each mutation was mapped onto this normalized scale using a linear interpolation approach: since repeat sequences are shorter than 100 nucleotides, such interpolation often caused a scenario where a mutation’s position did not correspond precisely to a single bin. The mutation position was then proportionally distributed across the two nearest bins based on its relative position. This normalization ensured a consistent and comparable framework for evaluating mutation distributions across different CRISPR subtypes, revealing distinct patterns in repeat sequence variability (see Figure 2). In a completely neutral mutation profile, one would expect that the number of mutations increases towards the end of the last repeat as mutations are less often eradicated there by spacer-repeat deletions and can thus accumulate over time. In a neutral repeat mutation process, a mutational hotspot within the repeat would be unlikely. Our analysis thus revealed that mutation positions within the repeat sequence vary distinctly across different CRISPR subtypes, indicating subtype-specific patterns of sequence variability and potential evolutionary pressures acting on repeat conservation.

**Figure 2.**
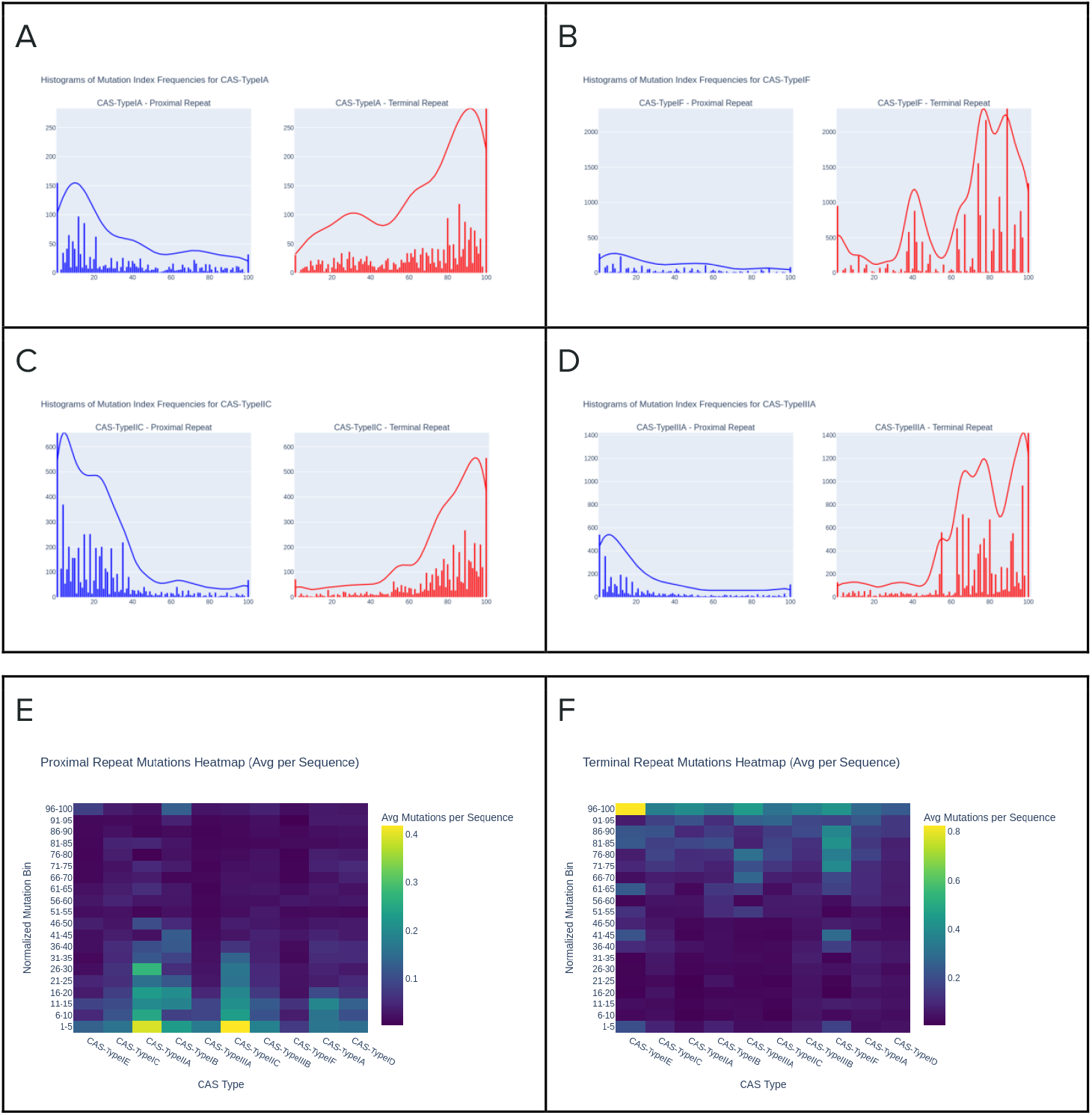
Mutation position distribution. Panels A-D show histograms of the mutation position normalized to 100 with the corresponding density curve. Different CRISPR subtypes clearly show different distributions of the mutation position. Panels E and F show heatmaps for proximal and terminal repeat mutations across all studied CRISPR subtypes.

### Frequency of Non-mutated proximal/terminal repeats is different for different CRISPR subtypes

To assess the prevalence of mutations in the leading (proximal) and terminal repeats, we conducted a comparative analysis of mutated versus non-mutated CRISPR arrays across different CRISPR subtypes. For each subtype, we identified the proportion of arrays in which both the leader-proximal and terminal repeats remained intact (without mutations) and compared it to the proportion of arrays exhibiting mutations in these regions (See Figure 3). This analysis allowed us to determine whether certain CRISPR subtypes are more prone to repeat sequence conservation or variability, providing insights into potential differences in the evolutionary constraints acting on the repeat sequences at the termini of CRISPR arrays (See Figure 3).

**Figure 3.**
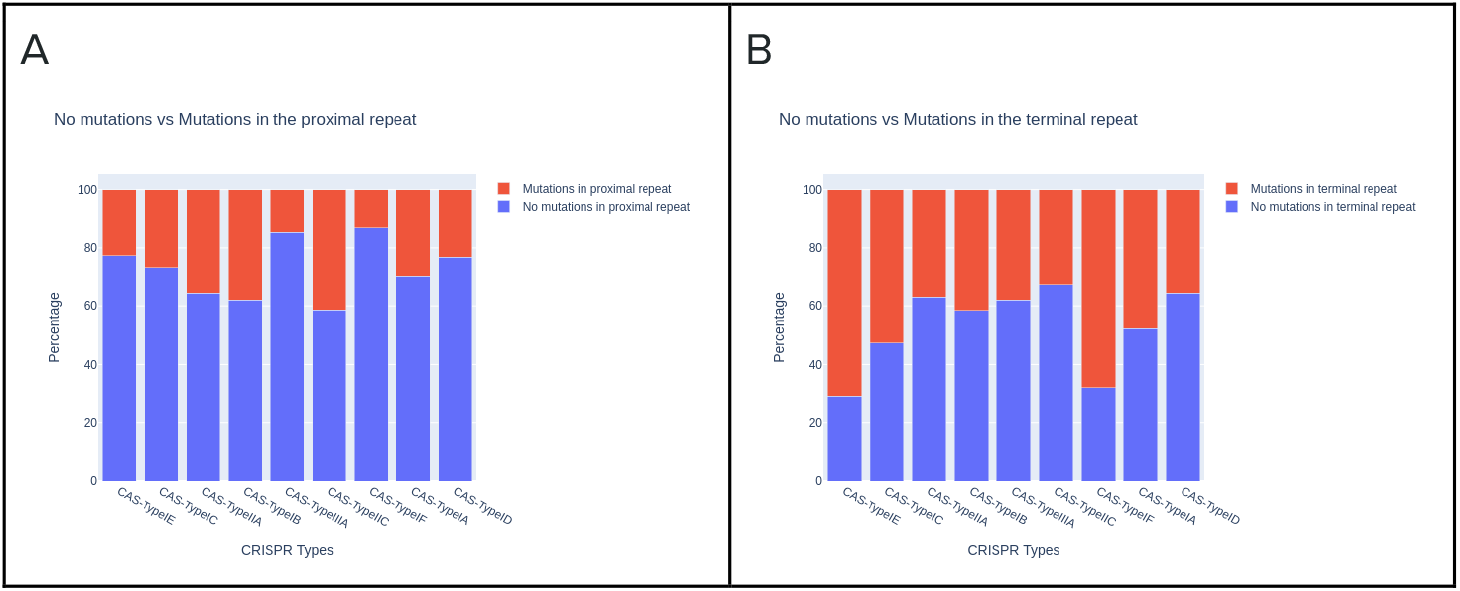
Distribution of mutated proximal and terminal repeat sequences across different CRISPR subtypes. The figure shows a clear difference in the quantity of mutated proximal repeat sequences shown in panel A from terminal repeat sequences shown in panel B. The percentage of mutated repeat sequences also varies between different CRISPR subtypes.

## Discussion

Our findings demonstrate different non-uniform distributions of repeat mutations across various CRISPR subtypes, with a notable increase in mutation frequencies observed at both the terminal and proximal ends of CRISPR arrays. This pattern indicates that repeat sequences may be influenced by unique evolutionary and mechanistic pressures, a topic that has received limited attention in prior investigations. By employing a large-scale dataset and computational analysis, we thoroughly analyse how repeat mutations affect the structural and functional evolution of CRISPR systems.

Previous research has primarily concentrated on the conservation of CRISPR repeat sequences, highlighting their crucial role in sustaining the stability and functionality of CRISPR arrays (Kunin, Sorek and Hugenholtz, 2007; Makarova et al., 2011). Nevertheless, our findings indicate that the accumulation of mutations occurs more frequently in specific regions of the array, particularly at its termini. This observation is consistent with earlier studies that propose that sequence variation within CRISPR loci may enhance bacterial adaptability by modulating the efficiency of spacer acquisition and the maturation of crRNA (Alkhnbashi et al. 2016, Alkhnbashi et al. 2014; Shah et al., 2013; Lange et al., 2013; Kunin et al., 2007). The subtype-specific variations in repeat mutation numbers further imply that certain CRISPR-Cas systems may prioritize stability, while others permit increased mutational flexibility. For instance, Type II systems, which depend on a well-defined repeat sequence for effective Cas9-mediated interference, demonstrate fewer mutations in comparison to Type I systems, where Cascade-mediated interference is potentially more accommodating of variations in repeat sequences (Van der Oost et al., 2014). This supports the hypothesis that the functionality and evolutionary path of a particular CRISPR-Cas system significantly influence the extent of repeat sequence conservation (Makarova et al., 2020).

Our findings indicate that repeat mutations may have a functional role in the processing and adaptation of CRISPR arrays. The observed increase in mutation frequency at the termini of arrays could be associated with the mechanisms involved in the acquisition of new spacers, which predominantly occur at the leader-proximal end (Barrangou et al., 2007). Should mutations in this region alter the recognition sites for the spacer integration machinery, they may influence the efficiency of new spacer acquisition and, consequently, the adaptive immunity of the host (Deltcheva et al., 2011). Furthermore, variations within repeat sequences could impact the stability and processing of crRNA, which is essential for effective interference activity. For instance, in Type III CRISPR-Cas systems, it has been demonstrated that variation in repeat sequences can affect the efficiency of RNA-targeting activity (Hale et al., 2009). Our research suggests that different CRISPR subtypes may exhibit differing levels of tolerance to repeat mutations, which could, in turn, influence their capacity to recognize and process foreign nucleic acids.

The observed variations in repeat mutations among CRISPR subtypes lend support to the hypothesis that these systems have developed distinct strategies to maintain their functionalities while permitting a certain degree of sequence variation. Type I and III systems, characterized by multi-protein complexes for interference, may demonstrate a greater tolerance for repeat mutations. In contrast, Type II systems, which depend on single-protein effectors such as Cas9, may exhibit more stringent requirements for sequence conservation (Charpentier et al., 2015). Notably, our data reveal that subtypes II-A and I-A show the most significant levels of discordance in strand direction prediction. This observation may suggest that these subtypes experience elevated repeat sequence evolution, potentially attributable to variations in their adaptation mechanisms (Brouns et al., 2008). A comprehensive understanding of these molecular-level differences could yield valuable insights into the diversification of CRISPR-Cas systems across bacterial and archaeal species.

CRISPR arrays undergo a dynamic process characterized by the acquisition and deletion of spacers, which is crucial for bacterial immunity. The research conducted by Fehrenbach et al. 2024 revealed that the frequency of spacer deletions is 374 times greater than that of mutations within the array, with these deletions predominantly occurring in clusters. This observation supports the hypothesis that the misalignment of repeats serves as a significant driver of spacer loss. Our findings regarding repeat mutations at the termini of CRISPR arrays suggest a potential correlation between variability in repeat sequences and the efficiency of spacer acquisition or deletion. It is noteworthy that new spacers are primarily integrated at the leader end of the array; thus, mutations in repeats in this region may impact the recognition sites for the adaptation machinery. This could influence both the efficiency of new spacer integration and the overall stability of the array. Furthermore, Fehrenbach et al. 2024 identified a decrease in spacer deletion frequencies at both ends of CRISPR arrays. This pattern mirrors our findings, which indicate elevated occurrences of repeat mutations at terminal repeats. Such observations imply an interconnectedness between the evolutionary dynamics of repeat mutations and the retention or loss of spacers, possibly driven by selection pressures that promote array stability while allowing for functional flexibility.

A comprehensive understanding of repeat mutation dynamics may enhance the design of synthetic CRISPR systems by increasing stability, thus reducing off-target effects and improving editing precision. Furthermore, insights into subtype-specific repeat flexibility could aid in the selection or engineering of Cas systems tailored explicitly for industrial or therapeutic applications.

## Conclusion

This study presents a thorough analysis of mutation dynamics within CRISPR array repeat sequences across diverse bacterial and archaeal genomes. Utilizing advanced bioinformatics tools such as CRISPRidentify, CRISPRcasFinder, and CRISPRcasIdentifier, we found a marked mutational bias at terminal repeats, exhibiting 5–7 fold higher mutation rates compared to internal repeats, particularly pronounced in subtype I-F (68.39 mutations per 1000 nucleotides).

These findings emphasize the evolutionary and functional significance of repeat mutations in balancing CRISPR array stability with adaptability. The subtype-specific mutation patterns uncovered hold significant practical implications, informing the development of more precise CRISPR-based genome-editing technologies and stable bacterial strains for industrial applications. Future studies should experimentally verify the functional consequences of repeat mutations to further harness their potential in synthetic biology.

## References

Abdelaal, A.S. and Yazdani, S.S. (2020) ‘Development and use of CRISPR in industrial applications’, in Genome Engineering via CRISPR-Cas9 System. Elsevier, pp. 177–197.

Alkhnbashi, O.S. et al. (2014) ‘CRISPRstrand: predicting repeat orientations to determine the crRNA-encoding strand at CRISPR loci’, Bioinformatics s, 30(17), pp. i489–96.

Alkhnbashi, O.S. et al. (2016) ‘Characterizing leader sequences of CRISPR loci’, Bioinformatics, 32(17), pp. i576–i585.

Alkhnbashi, O.S. et al. (2021) ‘CRISPRloci: comprehensive and accurate annotation of CRISPR-Cas systems’, Nucleic acids research, 49(W1), pp. W125–W130.

Barrangou, R. and Horvath, P. (2017) ‘A decade of discovery: CRISPR functions and applications’, Nature microbiology, 2(7), p. 17092.

Brouns, S.J.J. et al. (2008) ‘Small CRISPR RNAs guide antiviral defense in prokaryotes’, Science (New York, N.Y.), 321(5891), pp. 960–964.

Charpentier, E. et al. (2015) ‘Biogenesis pathways of RNA guides in archaeal and bacterial CRISPR-Cas adaptive immunity’, FEMS microbiology reviews, 39(3), pp. 428–441.

Couvin, D. et al. (2018) ‘CRISPRCasFinder, an update of CRISRFinder, includes a portable version, enhanced performance and integrates search for Cas proteins’, Nucleic acids research, 46(W1), pp. W246–W251.

Deltcheva, E. et al. (2011) ‘CRISPR RNA maturation by trans-encoded small RNA and host factor RNase III’, Nature, 471(7340), pp. 602–607.

Díez-Villaseñor, C. et al. (2013) ‘CRISPR-spacer integration reporter plasmids reveal distinct genuine acquisition specificities among CRISPR-Cas I-E variants of Escherichia coli’, RNA biology, 10(5), pp. 792–802.

Erdmann, S. and Garrett, R.A. (2012) ‘Selective and hyperactive uptake of foreign DNA by adaptive immune systems of an archaeon via two distinct mechanisms’, Molecular microbiology, 85(6), pp. 1044–1056.

Fehrenbach, A. et al. (2024) ‘SpacerPlacer: ancestral reconstruction of CRISPR arrays reveals the evolutionary dynamics of spacer deletions’, Nucleic acids research, 52(18), pp. 10862–10878.

de la Fuente-Núñez, C. and Lu, T.K. (2017) ‘CRISPR-Cas9 technology: applications in genome engineering, development of sequence-specific antimicrobials, and future prospects’, Integrative biology: quantitative biosciences from nano to macro, 9(2), pp. 109– 122.

Garrett, S.C. (2021) ‘Pruning and tending immune memories: Spacer dynamics in the CRISPR array’, Frontiers in microbiology, 12, p. 664299.

Hale, C.R. et al. (2009) ‘RNA-guided RNA cleavage by a CRISPR RNA-Cas protein complex’, Cell, 139(5), pp. 945–956.

Hale, C.R. et al. (2014) ‘Target RNA capture and cleavage by the Cmr type III-B CRISPR-Cas effector complex’, Genes & development, 28(21), pp. 2432–2443.

Hsu, P.D., Lander, E.S. and Zhang, F. (2014) ‘Development and applications of CRISPR-Cas9 for genome engineering’, Cell, 157(6), pp. 1262–1278.

Jinek, M. et al. (2012) ‘A programmable dual-RNA-guided DNA endonuclease in adaptive bacterial immunity’, Science (New York, N.Y.), 337(6096), pp. 816–821.

Koonin, E.V. and Makarova, K.S. (2017) ‘Discovery of Oligonucleotide Signaling Mediated by CRISPR-Associated Polymerases Solves Two Puzzles but Leaves an Enigma’, ACS chemical biology [Preprint]. Available at: 10.1021/acschembio.7b00713.

Koonin, E.V. and Makarova, K.S. (2019) ‘Origins and evolution of CRISPR-Cas systems’, Philosophical transactions of the Royal Society of London. Series B, Biological sciences, 374(1772), p. 20180087.

Kunin, V., Sorek, R. and Hugenholtz, P. (2007) ‘Evolutionary conservation of sequence and secondary structures in CRISPR repeats’, Genome biology, 8(4), p. R61.

Lange, S.J. et al. (2013) ‘CRISPRmap: an automated classification of repeat conservation in prokaryotic adaptive immune systems’, Nucleic acids research, 41(17), pp. 8034–8044.

Liao, C. et al. (2022) ‘Spacer prioritization in CRISPR-Cas9 immunity is enabled by the leader RNA’, Nature microbiology, 7(4), pp. 530–541.

Maier, L.-K. et al. (2017) ‘CRISPR and Salty: CRISPR-Cas Systems in Haloarchaea’, in RNA Metabolism and Gene Expression in Archaea. Cham: Springer International Publishing (Nucleic acids and molecular biology), pp. 243–269.

Makarova, K.S. et al. (2011) ‘Evolution and classification of the CRISPR-Cas systems’, Nature reviews. Microbiology, 9(6), pp. 467–477.

Makarova, K.S. et al. (2015) ‘An updated evolutionary classification of CRISPR-Cas systems’, Nature reviews. Microbiology, 13(11), pp. 722–736.

Makarova, K.S., Karamycheva, S., et al. (2019) ‘Predicted highly derived class 1 CRISPR-Cas system in Haloarchaea containing diverged Cas5 and Cas7 homologs but no CRISPR array’, FEMS microbiology letters, 366(7). Available at: 10.1093/femsle/fnz079.

Makarova, K.S., Gao, L., et al. (2019) ‘Unexpected connections between type VI-B CRISPR-Cas systems, bacterial natural competence, ubiquitin signaling network and DNA modification through a distinct family of membrane proteins’, FEMS microbiology letters, 366(8). Available at: 10.1093/femsle/fnz088.

Makarova, K.S. et al. (2020) ‘Evolutionary and functional classification of the CARF domain superfamily, key sensors in prokaryotic antivirus defense’, Nucleic acids research, 48(16), pp. 8828–8847.

Mitrofanov, A. et al. (2021) ‘CRISPRidentify: identification of CRISPR arrays using machine learning approach’, Nucleic acids research, 49(4), p. e20.

Naeem, M. and Alkhnbashi, O.S. (2023) ‘Current Bioinformatics Tools to Optimize CRISPR/Cas9 Experiments to Reduce Off-Target Effects’, International journal of molecular sciences, 24(7). Available at: 10.3390/ijms24076261.

Naito, Y. et al. (2015) ‘CRISPRdirect: software for designing CRISPR/Cas guide RNA with reduced off-target sites’, Bioinformatics (Oxford, England), 31(7), pp. 1120–1123.

Nickel, L. et al. (2019) ‘Cross-cleavage activity of Cas6b in crRNA processing of two different CRISPR-Cas systems in Methanosarcina mazei Gö1’, RNA biology, 16(4), pp. 492–503.

van der Oost, J. et al. (2014) ‘Unravelling the structural and mechanistic basis of CRISPR-Cas systems’, Nature reviews. Microbiology, 12(7), pp. 479–492.

Padilha, V.A. et al. (2020) ‘CRISPRcasIdentifier: Machine learning for accurate identification and classification of CRISPR-Cas systems’, GigaScience, 9(6). Available at: 10.1093/gigascience/giaa062.

Reimann, V. et al. (2017) ‘Structural constraints and enzymatic promiscuity in the Cas6-dependent generation of crRNAs’, Nucleic acids research, 45(2), pp. 915–925.

Shah, S.A. et al. (2019) ‘Comprehensive search for accessory proteins encoded with archaeal and bacterial type III CRISPR-cas gene cassettes reveals 39 new cas gene families’, RNA biology, 16(4), pp. 530–542.

Staals, R.H.J. et al. (2013) ‘Structure and activity of the RNA-targeting Type III-B CRISPR-Cas complex of Thermus thermophilus’, Molecular cell, 52(1), pp. 135–145.

Staals, R.H.J. et al. (2016) ‘Interference-driven spacer acquisition is dominant over naive and primed adaptation in a native CRISPR–Cas system’, Nature communications, 7(1), p. 12853.

Stachler, A.-E. et al. (2020) ‘Adaptation induced by self-targeting in a type I-B CRISPR-Cas system’, The Journal of biological chemistry, 295(39), pp. 13502–13515.

Touchon, M. and Rocha, E.P.C. (2010) ‘The small, slow and specialized CRISPR and anti-CRISPR of Escherichia and Salmonella’, PloS one, 5(6), p. e11126.

Vestergaard, G., Garrett, R.A. and Shah, S.A. (2014) ‘CRISPR adaptive immune systems of Archaea’, RNA biology, 11(2), pp. 156–167.

Westra, E.R. et al. (2012) ‘CRISPR immunity relies on the consecutive binding and degradation of negatively supercoiled invader DNA by Cascade and Cas3’, Molecular cell, 46(5), pp. 595–605.

Yosef, I., Goren, M.G. and Qimron, U. (2012) ‘Proteins and DNA elements essential for the CRISPR adaptation process in Escherichia coli’, Nucleic acids research, 40(12), pp. 5569– 5576.

